# Evidence for the role of cell reprogramming in naturally occurring cardiac repair

**DOI:** 10.1101/2020.10.16.342493

**Authors:** Nataliia V. Shults, Yuichiro J. Suzuki

## Abstract

Pulmonary arterial hypertension (PAH) is a fatal disease without a cure. If untreated, increased pulmonary vascular resistance kills patients within several years due to right heart failure. Even with the currently available therapies, survival durations remain short. By the time patients are diagnosed with this disease, the damage to the right ventricle (RV) has already developed. Therefore, agents that repair the damaged RV have therapeutic potential. We previously reported that cardiac fibrosis that occurs in the RV of adult Sprague-Dawley rats with PAH could naturally be reversed. We herein investigated the mechanism of this remarkable cardiac repair process. Counting of cardiomyocytes showed that the elimination of cardiac fibrosis is associated with the increased RV myocyte number, suggesting that new cardiomyocytes were generated. Immunohistochemistry showed the expression of α-smooth muscle actin and Sox-2 in RV myocytes of rats with PAH. Transmission electron microscopy detected the structure that resembles maturing cardiomyocytes in both the RV of PAH rats and cultured cardiomyocytes derived from induced pluripotent stem cells. We propose that the damaged RV in PAH can be repaired by activating the cell reprogramming mechanism that converts resident cardiac fibroblasts into induced cardiomyocytes.

## 1. Introduction

Pulmonary arterial hypertension (PAH) affects males and females of any age, including children. Despite the availability of approved drugs, PAH remains a fatal disease without a cure [1, 2]. The major pathogenic features that increase the pulmonary vascular resistance in PAH include the vasoconstriction and the development of vascular remodeling, in which pulmonary artery (PA) walls are thickened and the lumens are narrowed or occluded. Increased resistance puts strain on the right ventricle (RV), and right heart failure is the major cause of death among PAH patients [3, 4]. The median overall survival for patients diagnosed with PAH is 2.8 years from the time of diagnosis (3-year survival: 48%) if untreated [5, 6], Even with currently available therapies, the prognosis remains poor with only 58-75% of PAH patients surviving for 3 years [7–10]. PAH is a progressive disease; and by the time patients are diagnosed, RV damage has often already occurred.

RV failure is the major cause of death among patients with PAH. However, no treatment strategies are available to manage the RV dysfunctions. Physiologically, the RV needs to cope with fold changes in PA pressure. Thus, the RV is capable of adapting to increased pressure. Similarly to human patients with PAH, we found that the RV of Sprague-Dawley (SD) rats treated with SU5416 and hypoxia to produce PAH suffer from severe cardiac fibrosis at 8 to 17 weeks after the initiation of the SU5416/hypoxia treatment [11]. Remarkably, at 35 weeks after the initiation of the SU5416/hypoxia treatment, RV fibrosis was found to be resolved in these rats, despite the RV pressure remained high [12]. Thus, the RV remodeling can naturally be reversed in these animals, providing an interesting model of RV repair. Understanding the mechanism of such naturally occurring events should shed a lighten developing therapeutic strategies to repair the cardiac damage in human patients. The present study examined the mechanism of this cardiac repair process.

## 2. Materials and Methods

### Experimental animals

Male adult SD rats (Charles River Laboratories International, Inc., Wilmington, MA, USA) were subcutaneously injected with SU5416 (20 mg/kg body weight; MedChem Express, Monmouth Junction, NJ, USA), maintained in hypoxia for 3 weeks [11, 12] and then in normoxia for up to 32 weeks (35-week time points). Animals were subjected to hypoxia in a chamber (30”w x 20”d x 20”h) regulated by an OxyCycler Oxygen Profile Controller (Model A84XOV; BioSpherix, Redfield, NY, USA) set to maintain 10% O_2_ with an influx of N_2_ gas, located in the animal care facility at the Georgetown University Medical Center [11, 12]. Ventilation to the outside of the chamber was adjusted to remove CO_2_, such that its level did not exceed 5,000 ppm. Control animals were subjected to ambient 21% O_2_ (normoxia) in another chamber. Animals were fed normal rat chow. Animals were anesthetized and euthanized by excising the heart and the lungs.

The Georgetown University Animal Care and Use Committee approved all animal experiments, and the investigation conformed to the National Institutes of Health (NIH) Guide for the Care and Use of Laboratory Animals.

### Immunohistochemistry (IHC)

RV tissues were immersed in buffered 10% formalin at room temperature, and were embedded in paraffin. These paraffin-embedded tissues were cut and mounted on glass slides. IHC was performed using horseradish peroxidase (HRP) labeled polymer and 3,3’-diaminobenzidine (DAB) chromagen (Agilent Technologies, Santa Clara, CA, USA) with α-smooth muscle actin (αSMA; Catalog # ab32575) and Sox2 (Catalog # ab97959) antibodies (Abeam, Cambridge, UK).

### Transmission electron microscopy (TEM)

The RV free wall tissues of rats subjected to SU5416/hypoxia as well as cultured cardiomyocytes derived from human induced pluripotent stem cells (iPSCs) purchased from Cell Applications, Inc. (San Diego, CA) were fixed in the 2.5% glutaraldehyde/0.05M cacodylate solution, post-fixed with 1% osmium tetroxide and embedded in EmBed812. Ultrathin sections (70 nm) were poststained with uranyl acetate and lead citrate and examined in the Talos F200X FEG transmission electron microscope (FEI, Hillsboro, OR, USA) at 80 KV located at the George Washington University Nanofabrication and Imaging Center. Digital electron micrographs were recorded with the TIA software (FEI).

### Statistical analysis

Means and standard errors were calculated. Comparisons between three groups were analyzed by using one-way analysis of variance (ANOVA) with a Student-Newman-Keuls post-hoc test using the GraphPad Prism (GraphPad Software, Inc., La Jolla, CA, USA). P< 0.05 was considered to be significant.

## 3. Results

Since the initial discovery that treating SD rats with the SU5416 injection plus chronic hypoxia promoted severe PAH with pulmonary vascular lesions that resemble those of human patients [13], this experimental model has become a gold standard in the research field of PAH. Experimental design usually involves the single subcutaneous injection of SU5416, followed by subjecting to chronic hypoxia for 3 weeks. In many studies, rats are then maintained in normoxia for 2 to 5 weeks, and PAH and pulmonary vascular remodeling are observed [13–17]. At this stage, some laboratories including ours have reported that the RV is severely damaged with fibrosis [11, 14, 18, 19]. At 8 to 17 weeks after the SU5416 injection, however, we found that, despite the occurrence of severe fibrosis, the RV contractility is maintained or even improved, perhaps due to the formation of ‘super’ RV myocytes [11]. Moreover, our more recent results demonstrated that, at 35 weeks (3 weeks hypoxia followed by 32 weeks of normoxia), RV fibrosis was largely resolved, despite the RV pressure remained high [12]. The number of myofibroblasts that contribute to the formation of fibrosis, as detected using a well utilized marker αSMA, is increased in non-vessel regions of the RV of SD rats with severe PAH as well as RV fibrosis, and this expression declined in rats with repaired RV at 35 week after the SU5416 injection [12]. Thus, the SU5416/hypoxia treatment does not cause the death ofSD rats and the damaged RV can be repaired naturally. Since RV myocytes were not hypertrophied at 35-weeks, the number of myocytes must have been increased to fill the post-fibrotic areas. Indeed counting cardiomyocytes indicated that, compared to the 17-week time point when fibrosis was present, the number of cardiomyocytes increased at 35-weeks when the RV was repaired (Figure 1).

**Fig. 1:**
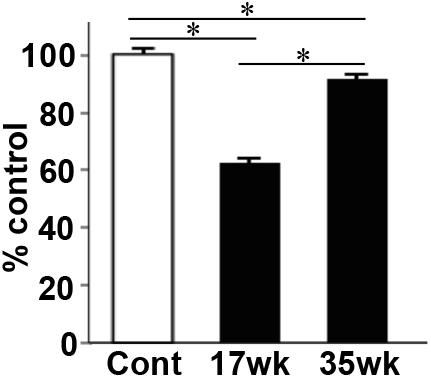
Restoration of RV cardiomyocytes. SU54l6-injected SD rats were subjected to 3 weeks hypoxia and then maintained in normoxia to promote PAH. 17 and 35 weeks after the SU5416 injection, RV myocardium tissues were fixed in formalin, embedded in paraffin, and subjected to H&E staining. The number of cardiomyocytes were counted and expressed as % of the control rats (Cont). SD rats with PAH at 17 weeks after the initiation of SU5416/ hypoxia had significantly reduced RV myocyte number compared to healthy controls. This decrease in RV myocyte number was restored at 35 weeks. The symbol * denotes that the values are significantly different from each other at *p*<0.05.

Based on these results, we hypothesize that these SD rats possess a mechanism that repairs the damaged heart in response to PAH. This raises a question on where regenerated cardiomyocytes come from when the RV repair occurs. It is now known that some adult cardiomyocytes are capable of proliferating [20]. However, the cardiac renewal through the cardiomyocyte proliferation would be too slow to replace large fibrotic areas that were seen in our experimental model. Another possibility is that new cardiomyocytes were regenerated from cardiac progenitor cells. We have identified c-kit and isl1-positive cardiac progenitor cells in the RV of SD rats, however, their levels were not altered in PAH rats (data not shown).

Our previous experiments described in Zungu-Edmondson et al. [12] using the αSMA antibody were originally performed for the purpose of detecting myofibroblasts, which express σ,SMA. By examining αSMA IHC slides from RVs of SD rats with PAH (at 17-weeks) with a larger magnification (x1,000) as shown in Figure 6A of Zungu-Edmondson et al. [12], we noticed that some brown αSMA stains, in addition to myofibroblasts indicated by arrows, also occurred on cardiomyocytes. We initially discarded these observations by thinking that they are non-specific artifacts. However, further examinations of a number of IHC slides made us convinced that cardiomyocytes are indeed stained with the αSMA antibody. This seems to occur regionally as a group in the myocardial walls (Figure 2*A*). Figure 2B shows the amplified view of Figure 2A clearly demonstrating that these cardiomyocytes with clear striations express αSMA. By contrast, control RVs from healthy rats without PAH did not exhibit the staining of cardiomyocytes with the αSMA antibody (Figure 2C).

**Fig. 2:**
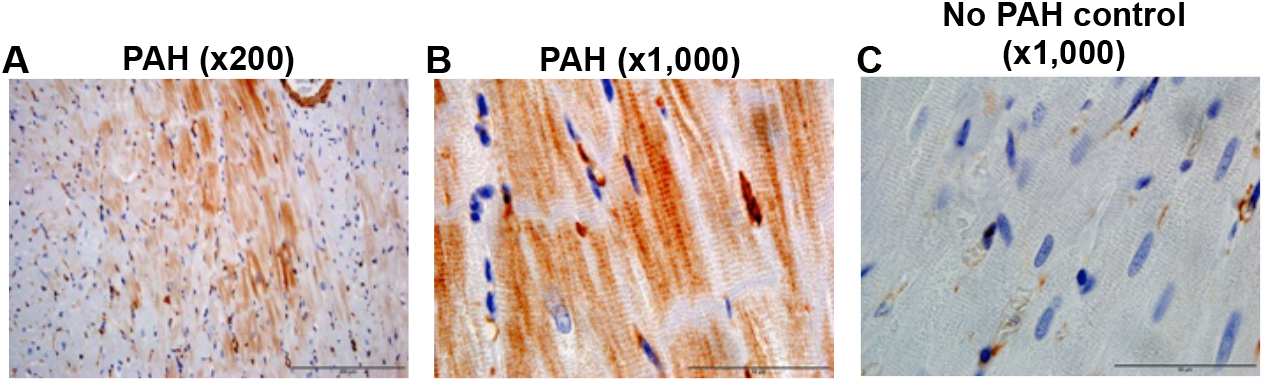
Discovery of naturally-occurring induced cardiomyocytes (iCMs)? SU5416- injected SD rats were subjected to 3 weeks hypoxia and then maintained in normoxia to promote PAH. 17 weeks after the injection, myocardium tissues were fixed in formalin, embedded in paraffin, and subjected to IHC using the antibody against αSMA (brown stains). (A and B) αSMA IHC results shown at x200 and x1,000 magnifications, which clearly show the brown stains in RV cardiomyocytes of SD rats with PAH. (D) This brown αSMA stain was not observed in control rat RVs without PAH. Scale bars indicate 200 μm for x200 and 50 μm for x1,000.

Cell reprogramming is defined as the conversion of one specific cell type to another. This technology has gained considerable attention when Prof. Shinya Yamanaka discovered the means to convert fibroblasts into iPSCs and received the Nobel Prize [21]. In their study, stem cell-related transcription factors including Oct4 and Sox2 were used to convert somatic cells to pluripotent cells that can be differentiated into various cell types including cardiomyocytes [22]. More recently, a combination of cardiac-specific transcription factors was found to directly convert fibroblasts into cardiomyocytes [23, 24]. Figure 3A shows our TEM study of cardiomyocytes derived from iPSCs. These induced cardiomyocytes (iCMs) are capable of beating, express contractile proteins, and exhibit an organized sarcomere structure with clear striations and Z-lines (Figure 3A). In addition, some regions with the not well-defined sarcomere organization, which could be in the process of maturing into iCMs were also identified in these TEM images (Figure 3B). iCMs have been shown to express αSMA [25], while normal adult cardiomyocytes do not. Thus, αSMA-positive cardiomyocytes we observed in adult SD rats with PAH could be iCM-like cells. Consistently with this hypothesis, our examination of TEM images revealed that the structure resembling cardiomyocytes maturing from iPSCs as observed in cultured cells (Figure 3B) also occurs in the RV of SD rats with PAH (Figure 4B).

**Fig. 3:**
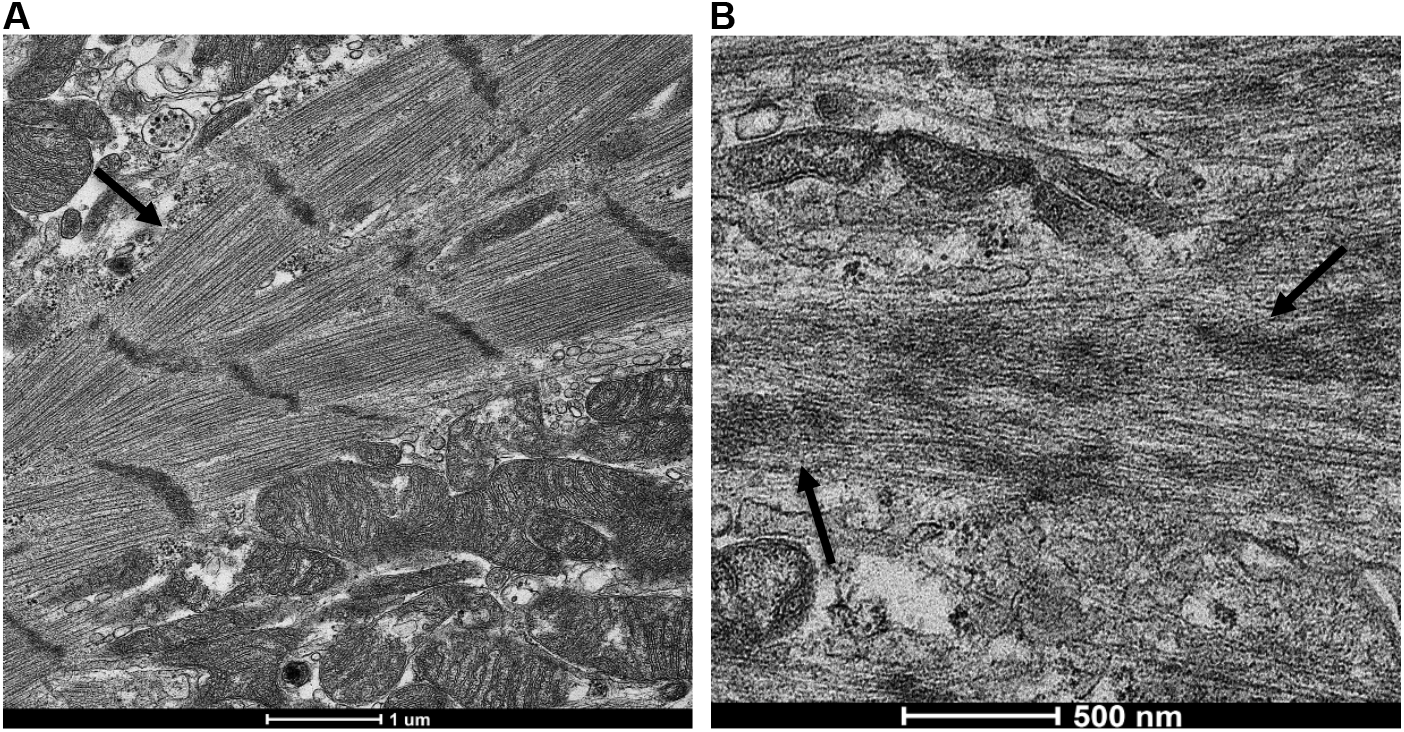
TEM image of cardiomyocytes derived from iPSCs. The reprogramming technology was used to convert cultured human fibroblasts into iPSCs, then to cardiomyocytes (Cell Applications, Inc.). Fixed iPSC-derived cardiomyocytes were observed under a transmission electron microscope. (A) The representative image shows cardiomyocytes with clear striations and sarcomere structures (arrow), indicating that iPSCs indeed can become matured cardiomyocytes. Magnification, x5,500. (B) We also identified some regions with the not well defined sarcomere organization (arrows), which may be in the process of maturing into cardiomyocytes. Magnification, x14,000.

**Fig. 4:**
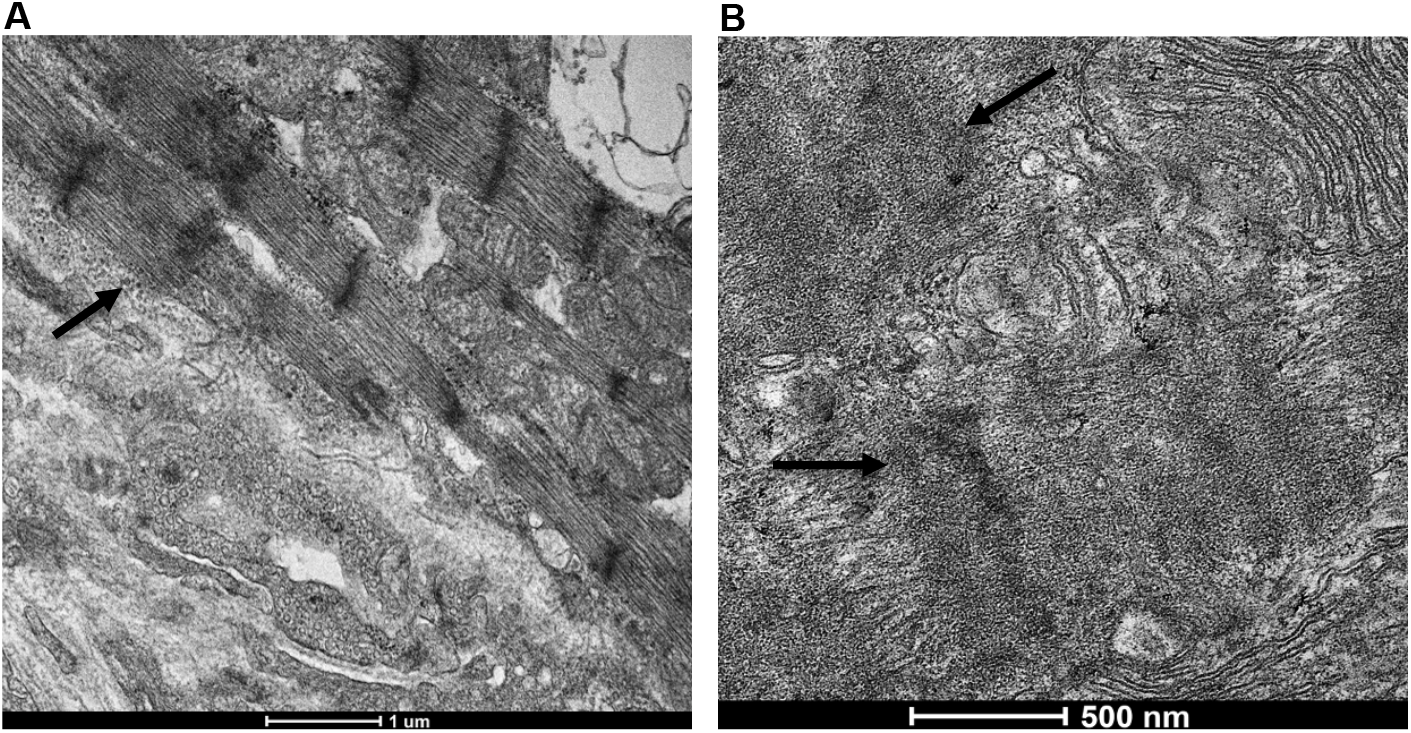
TEM identification of maturing ICM-like cells in the RV of PAH rats. SD rats were injected with SU5416, subjected to 3 weeks hypoxia, and maintained in normoxia. 20 weeks after the injection, RV tissues were fixed and analyzed by TEM. (A) The image shows normal cardiomyocytes with clear striations and sarcomere structures (arrow). Magnification, x5,500. (B) We also identified some regions with the not well defined sarcomere organization (arrows), which may be in the process of maturing into iCMs, similar to the structure shown in Fig. 3B. Magnification, x14,000.

These results would support the concept that RVs of SD rats with PAH have iCMs-like cells that are produced via cell reprogramming. However, the αSMA expression can be induced by the fetal gene program mechanism that is associated with cardiac hypertrophy, independent of cell reprogramming. Thus, we examined if factors more directly related to cell reprogramming are expressed in these cardiomyocytes. We found that cardiomyocytes of RV tissues from SD rats with PAH also express Sox2 (Figures 5A and 5B), a stem cell-related transcription factor that has been used to generate iPSCs [22]. By contrast, no Sox2 stains were observed in healthy control SD rats without PAH (Figure 5C).

**Fig. 5.**
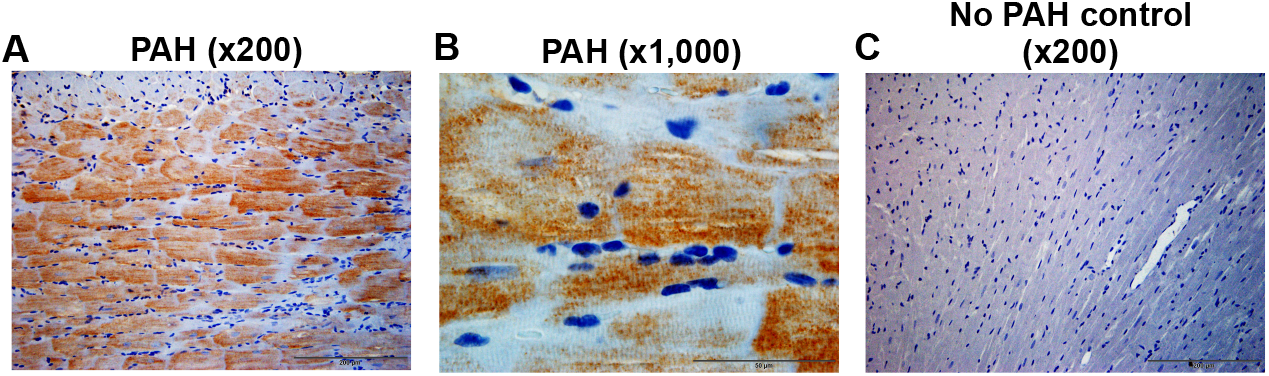
Expression of Sox2. SU5416-injected SD rats were subjected to 3 weeks hypoxia and then maintained in normoxia to promote PAH. 17 weeks after the injection, hearts were fixed and subjected to IHC with the antibody against Sox2 (brown stains). (A and B) Sox2 IHC results shown at x200 and at x1,000 magnifications, which clearly show the brown stains in RV cardiomyocytes of SD rats with PAH. (C) This brown Sox2 stain was not observed in control rat RVs without PAH. Scale bars indicate 200 μm for x200 and 50 μm for x1,000.

## 4. Discussion

In response to pressure overload, the heart ventricles undergo a series of adaptive events. In response to systemic and pulmonary hypertension, the left ventricle (LV) and the RV, respectively hypertrophy in order to increase the force of muscle contraction. Concentric hypertrophy is the first change that occurs in response to chronic pressure overload, and this compensatory mechanism allows for improved cardiac output. Exercise-induced cardiac hypertrophy, for example, increases the force of contraction in accordance the needs associated with strenuous exercise and training. This adaptive feature is reversible. However, in chronic disease conditions, this compensatory mechanism thickens the ventricular wall too much in a manner that decreases the stroke volume and thus the cardiac output. This results in the second adaptation to decrease the ventricular wall thickness. In case of the LV, the transition from concentric to eccentric hypertrophy predominates, resulting in the dilated LV with the thin ventricular wall. This event, however, does not seem to occur in the RV in response to chronic pulmonary hypertension and the hallmark of cor pulmonale is that the RV myocytes remain concentrically hypertrophied at the time of heart failure. In the RV, the attempt to decrease the RV wall thickness is expected to be related to the promotion of cell death, although it is unclear whether apoptosis really occurs in the RV of human patients with pulmonary hypertension [26]. However, apoptotic cardiomyocytes have been detected in the RV of a rat model of pulmonary hypertension [11]. In these rats, regions of the RV where cardiomyocytes die get filled with fibrosis.

Using well-studied model of PAH, in which SD rats are treated with the SU5416 injection and chronic hypoxia, we previously found that RVs were capable of maintaining sufficient force of muscle contraction even severe fibrosis occurred at 8 to 17 weeks after the initiation of the SU5416/hypoxia treatment [11]. We postulated that this is due to the formation of “super RV myocytes” that are capable of eliciting stronger force of contraction via a mechanism involving the downregulation of calsequestrin 2, the major Cambinding protein of the sarcoplasmic reticulum.

Further, remarkably at 35 weeks after the SU5416/hypoxia initiation, these fibrosis regions disappear and are filled with newly formed cardiomyocytes [12]. The present study indeed showed that the number of cardiomyocytes were increased at the 35-week time point, compared to the 17-week time point. This increased number of cardiomyocytes was found to be associated with the production of RV cardiomyocytes that express αSMA and Sox2 as well as the occurrence of the structure visible in TEM that resembles maturing iCMs similar to those observed in cultured cardiomyocytes derived from iPSCs.From these results, we hypothesize that the damaged RV due to PAH can naturally be repaired in SD rats through a mechanism that involves the conversion of resident cardiac fibroblasts into iCMs via cell reprogramming (Figure 6). Our finding also suggests that the nature possesses the ability for maturing iCMs into functional cardiomyocytes that are capable of eliciting strong muscle contraction through a “functionalization” process (Figure 6). Our results so far provided evidence to support this novel mechanism of cardiac regeneration, however, further work is needed to prove this concept.

**Fig. 6.**
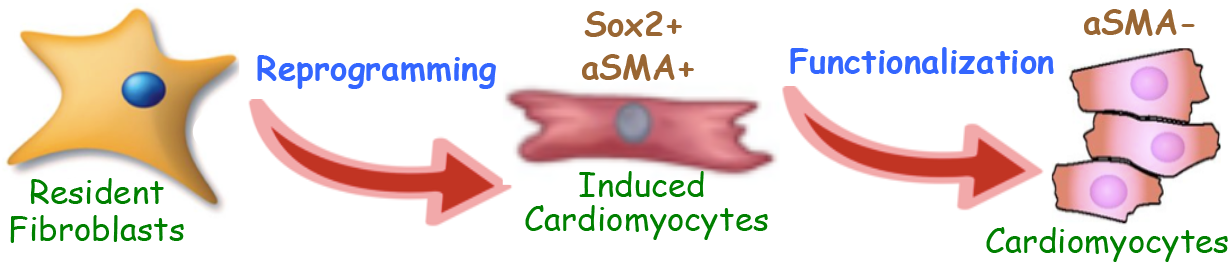
Scheme depicting our hypothesis. SD rats possess an RV repair mechanism, in which naturally occurring cell reprogramming process converts resident fibroblasts into αSMA-and Sox2-positive induced cardiomyocytes (iCMs). These rats also possess the functionalization mechanism that makes functional cardiomyocytes that can elicit strong muscle contraction.

Understanding of the endogenous means to repair the heart is important, not only to provide basic knowledge of cardiac physiology, but also to develop therapeutic strategies to treat conditions in which cardiomyocytes are damaged. Along with the cardiomyocyte proliferation and the involvement of cardiac progenitor cells, our results suggest a novel mechanism of cardiac regeneration through cell reprogramming. Defining this naturally occurring mechanism should contribute to the development of technologies to convert resident cardiac fibroblasts into cardiomyocytes to save lives.

## 5. Conclusion

The present study generated evidence to support a fascinating and novel mechanism of naturally occurring cardiac repair that should be important for the development of effective therapeutic strategies to treat cardiac failure. This novel mechanism involves the conversion of resident cardiac fibroblasts into iCMs through the cell reprogramming process that appears to share events occurring in iPSC biology.

## 6. Acknowledgments

This work was supported in part by the NIH (grant numbers RoiHL072844, R21AI142649, R03AG059554, and R03AA026516) to Y.J.S. The content is solely the responsibility of the authors and does not necessarily represent the official views of the NIH.

## 7. Conflict of Interest

None.

## Notes

### Competing Interest Statement

The authors have declared no competing interest.

## References

[1] Fallah F. Recent strategies in treatment of pulmonary arterial hypertension, a review. Glob J Health Sci. 2015;7:307–322.

[2] Rosenkranz S. Pulmonary hypertension 2015: current definitions, terminology, and novel treatment options. Clin Res Cardiol. 2015;104:197–207.

[3] Delcroix M, Naeije R. Optimising the management of pulmonary arterial hypertension patients: emergency treatments. Eur Respir Rev. 2010;19:204–211.

[4] McLaughlin VV, Shah SJ, Souza R, Humbert M. Management of pulmonary arterial hypertension. J Am Coll Cardiol. 2015;65:1976–97.

[5] D Alonzo GE, Barst RJ, Ayres SM, Bergofsky EH, Brundage BH, Detre KM, Fishman AP, Goldring RM, Groves BM, Kernis JT, et al. Survival in patients with primary pulmonary hypertension. Results from a national prospective registry. Ann Intern Med. 1991; 115:343–349.

[6] Runo JR, Loyd JE. Primary pulmonary hypertension. Lancet. 2003;361:1533–1544.

[7] Benza RL, Miller DP, Frost A, Barst RJ, Krichman AM, and McGoon MD. Analysis of the lung allocation score estimation of risk of death in patients with pulmonary arterial hypertension using data from the REVEAL Registry. Transplantation. 2010;90:298–305.

[8] Humbert M, Sitbon O, Yaïci A, Montani D, O’Callaghan DS, Jaïs X, Parent F, Savale L, Natali D, Günther S, Chaouat A, Chabot F, Cordier JF, Habib G, Gressin V, Jing ZC, Souza R, Simonneau G; French Pulmonary Arterial Hypertension Network. Survival in incident and prevalent cohorts of patients with pulmonary arterial hypertension. Eur Respir J. 2010;36:549–555.

[9] Thenappan T, Shah SJ, Rich S, Tian L, Archer SL, Gomberg-Maitland M. Survival in pulmonary arterial hypertension: a reappraisal of the NIH risk stratification equation. Eur Respir J. 2010;35:1079–1087.

[10] Olsson KM, Delcroix M, Ghofrani HA, Tiede H, Huscher D, Speich R, Grünig E, Staehler G, Rosenkranz S, Halank M, Held M, Lange TJ, Behr J, Klose H, Claussen M, Ewert R, Opitz CF, Vizza CD, Scelsi L, Vonk-Noordegraaf A, Kaemmerer H, Gibbs JS, Coghlan G, Pepke-Zaba J, Schulz U, Gorenflo M, Pittrow D, Hoeper MM. Anticoagulation and survival in pulmonary arterial hypertension: results from the Comparative, Prospective Registry of Newly Initiated Therapies for Pulmonary Hypertension (COMPERA). Circulation. 2014;129:57–65.

[11] Zungu-Edmondson M, Shults NV, Wong CM, Suzuki YJ. Modulators of right ventricular apoptosis and contractility in a rat model of pulmonary hypertension. Cardiovasc Res. 2016;110:30–39.

[12] Zungu-Edmondson M, Shults NV, Melnyk O, Suzuki YJ. Natural reversal of pulmonary vascular remodeling and right ventricular remodeling in SU54i6/hypoxia-treated Sprague-Dawley rats. PLoS One. 2017;12:e0182551.

[13] Taraseviciene-Stewart L, Kasahara Y, Alger L, Hirth P, McMahon G, Waltenberger J, Voelkel NF, Tuder RM. Inhibition of the VEGF receptor 2 combined with chronic hypoxia causes cell death-dependent pulmonary endothelial cell proliferation and severe pulmonary hypertension. FASEB J. 2001;15:427–438.

[14] Oka M, Homma N, Taraseviciene-Stewart L, Morris KG, Kraskauskas D, Burns N, Voelkel NF, McMurtry IF. Rho kinase-mediated vasoconstriction is important in severe occlusive pulmonary arterial hypertension in rats. Circ Res. 2007; 100:923–929.

[15] Ibrahim YF, Wong CM, Pavlickova L, Liu L, TrasarL, Bansal G, Suzuki YJ. Mechanism of the susceptibility of remodeled pulmonary vessels to drug-induced cell killing. J Am Heart Assoc. 2014;3:e000520.

[16] Alzoubi A, Toba M, Abe K, O‘Neill KD, Rocic P, Fagan KA McMurtry IF, Oka M. Dehydroepiandrosterone restores right ventricular structure and function in rats with severe pulmonary arterial hypertension. Am J Physiol Heart Circ Physiol. 2013;304: H1708–H1718.

[17] Wang X, Ibrahim YF, Das D, Zungu-Edmondson M, Shults NV, Suzuki YJ. Caŗfilzomib reverses pulmonary arterial hypertension. Cardiovasc Res. 2016;110:188–199.

[18] Bogaard HJ, Natarajan R, Henderson SC, Long CS, Kraskauskas D, Smithson L, Ockaili R, McCord JM, Voelkel NF. Chronic pulmonary artery pressure elevation is insufficient to explain right heart failure. Circulation. 2009;120:1951–1960.

[19] Bogaard HJ, Natarajan R, Mizuno S, Abbate A, Chang PJ, Chau VQ, Hoke NN, Kraskauskas D, Kasper M, Salloum FN, Voelkel NF. Adrenergic receptor blockade reverses right heart remodeling and dysfunction in pulmonary hypertensive rats. Am J Respir Crit Care Med. 2010;182:652–660.

[20] Yuan X, Braun T. Multimodal regulation of cardiac myocyte proliferation. Circ Res. 2017;121:293–309.

[21] Yamanaka S. Induced pluripotent stem cells: Past, present, and future. Cell Stem Cell. 2012;10:678–684.

[22] Takahashi K, Yamanaka S. Induction of pluripotent stem cells from mouse embryonic and adult fibroblast cultures by defined factors. Cell. 2006;126:663–676.

[23] Ieda M, Fu JD, Delgado-Olguin P, Vedantham V, Hayashi Y, Bruneau BG, Srivastava D. Direct reprogramming of fibroblasts into functional cardiomyocytes by defined factors. Cell. 2010;142:375–386.

[24] Song K, Nam YJ, Luo X, Qi X, Tan W, Huang GN, Acharya A, Smith CL, Tallquist MD, Neilson EG, Hill JA, Bassel-Duby R, Olson EN. Heart repair by reprogramming non-myocytes with cardiac transcription factors. Nature. 2012;485:599–604.

[25] Cao N, Huang Y, Zheng J, Spencer CI, Zhang Y, Fu JD, Nie B, Xie M, Zhang M, Wang H, Ma T, Xu T, Shi G, Srivastava D, Ding S. Conversion of human fibroblasts into functional cardiomyocytes by small molecules. Science. 2016; 352:1216–1220.

[26] Voelkel NF, Quaife RA, Leinwand LA, Barst RJ, McGoon MD, Meldrum DR, Dupuis J, Long CS, Rubin LJ, Smart FW, Suzuki YJ, Gladwin M, Denholm EM, Gail DB; Right ventricular function and failure: report of a National Heart, Lung, and Blood Institute working group on cellular and molecular mechanisms of right heart failure. Circulation. 2006;114:1883–1891.

